# Agronomic Linked Data (AgroLD): a Knowledge-based System to Enable Integrative Biology in Agronomy

**DOI:** 10.1101/325423

**Authors:** Aravind Venkatesan, Gildas Tagny, Nordine El Hassouni, Imene Chentli, Valentin Guignon, Clement Jonquet, Manuel Ruiz, Pierre Larmande

## Abstract

Recent advances in high-throughput technologies have resulted in a tremendous increase in the amount of omics data produced in plant science. This increase, in conjunction with the heterogeneity and variability of the data, presents a major challenge to adopt an integrative research approach. We are facing an urgent need to effectively integrate and assimilate complementary datasets to understand the biological system as a whole. The Semantic Web offers technologies for the integration of heterogeneous data and their transformation into explicit knowledge thanks to ontologies. We have developed the Agronomic Linked Data (AgroLD – www.agrold.org), a knowledge-based system relying on Semantic Web technologies and exploiting standard domain ontologies, to integrate data about plant species of high interest for the plant science community e.g., rice, wheat, arabidopsis. We present some integration results of the project, which initially focused on genomics, proteomics and phenomics. AgroLD is now an RDF (Resource Description Format) knowledge base of 100M triples created by annotating and integrating more than 50 datasets coming from 10 data sources –such as Gramene.org and TropGeneDB– with 10 ontologies –such as the Gene Ontology and Plant Trait Ontology. Our evaluation results show users appreciate the multiple query modes which support different use cases. AgroLD’s objective is to offer a domain specific knowledge platform to solve complex biological and agronomical questions related to the implication of genes/proteins in, for instances, plant disease resistance or high yield traits. We expect the resolution of these questions to facilitate the formulation of new scientific hypotheses to be validated with a knowledge-oriented approach.

## Introduction and Background

Agronomy is a multi-disciplinary scientific discipline that includes research areas such as plant molecular biology, physiology and agro-ecology. Agronomic research aims to improve crop production and study the environmental impact on crops. Accordingly, researchers need to understand the implications and interactions of the various biological processes, by linking data at different scales (e.g., genomics, proteomics and phenomics). We are currently witnessing rapid advances in high throughput and information technologies that continue to drive a flood of data and analysis techniques within the domains mentioned above. However, much of these data or information are dispersed across different domain or model specific databases, varied formats and representations e.g., TAIR, GrainGenes and Gramene. Therefore, using these databases more effectively and adopting an integrative approach remains a major challenge.

Among the numerous research directions that the field of bioinformatics has taken, knowledge management has become a major area of research, focused on logically interlinking information and the representation of domain knowledge [1]. To this end, ontologies have become a cornerstone in the representation of biological and more recently agronomical knowledge [2]. Ontologies provide the necessary scaffold to represent and formalize biological concepts and their relationships. Currently, numerous applications exploit the advantages offered by biological ontologies such as: the Gene Ontology [3] –widely used to annotate genes and their products– Plant Ontology [4], Crop Ontology [5], Environment Ontology [6], to name a few. Ontologies have opened the space to various types of semantic applications [7,8] to data integration [9], and to decision support [10]. Semantic interoperability has been identified as a key issue for agronomy, and the use of ontologies declared a way to address it [11]. Furthermore, efficient knowledge management requires the adoption of effective data integration methodologies. This involves efficient semantic integration of the disparate data sources, making information machine-readable and interoperable. Accordingly, Semantic Web standards and technologies enforced by the W3C, and embracing Tim Berners-Lee’s vision [12], offers a solution to facilitate integration and interoperability of highly diverse and distributed data resources. The Semantic Web technologies stack includes among others the following W3C Recommendations: the Resource Description Framework (RDF) [13] as a backbone language to describe resources with triples, RDF Schema (RDFS) [14] to build lightweight data schemas, Web Ontology Language (OWL) [15] to build semantically rich ontologies and the SPARQL Query Language (SPARQL) [16] to query RDF data. All of the previous languages rely on Unique Resource Identifiers (URIs) to define a resource and its components, enabling data interoperability across the Web. RDF describes a resource and its relationships/properties in the form of simple triples, i.e., *Subject-Predicate-Object* offering a very convenient framework for integrating data across multiple platforms assuming the platforms share some common vocabularies to describe their objects. These triples can be combined to construct large networks of information (also known as RDF graphs). A successfully implemented Semantic Web application allows scientists to pose very complex questions through a query or a set of queries that would return highly relevant answers to those questions, facilitating the formulation of research hypotheses [17,18].

There are other approaches to meet the current data integration challenges, e.g., data warehouses. For instance, Intermine [19] has developed a sophisticated application to accommodate the dynamic nature of biological data and simplify data integration. However, with integrative biology gaining popularity, it is necessary to preserve and share the semantics between the various datasets and make information machine interoperable, enabling large scale analyses of information available over the Web. The Semantic Web approach provides an added value, playing a complementary role to the traditional methods of data integration.

In the recent years, the biomedical community has strongly embraced the Semantic Web vision as demonstrated by a number of initiatives to provide ontologies [20,21] and use them for producing semantically rich data such as in Bio2RDF [22], OpenPHACTS [23], Linked Life Data [24], KUPKB [25], and the EBI RDF Platform [26]. In particular, OpenPHACTS serves as a good example of what can be achieved by using Semantic Web knowledge bases. The OpenPHACTS Explorer (http://www.openphacts.org/open-phacts-discovery-platform/explorer) provides use case driven tools that aid in browsing and visualizing the underlying knowledge represented in RDF which is very convenient for biologists.

Currently, there is a growing awareness within the agronomic domain towards efficient data interoperability and integration [2,27,28]. The need for an umbrella approach for providing uniform data is a widely-discussed topic. For instance, the Agriculture Data Interoperability Interest Group (https://rd-alliance.org/groups/agriculture-data-interest-group-igad.html) instituted by the Research Data Alliance (RDA) and agINFRA EU project (www.aginfra.eu) are initiatives that work on improving data standards and promoting data interoperability in agriculture. Moreover, the community has recently also started to adopt AgroPortal [11] as an vocabulary and ontology repository for agronomy –and related domains such as nutrition, plant sciences and biodiversity– that support browsing, searching and visualizing domain relevant ontologies, ontology alignments and creation of semantic annotations. While plant-centric ontologies are now being used to annotate data by various databases developers [2,5,28], unlike in the biomedical domain, the adoption of Semantic Web in agronomy is yet to be completely exploited. Given that agronomic studies involve multiple domains, publicly available knowledge bases such as EBI RDF, Linked Life Data and Bio2RDF serves only limited agronomical information. Hence, it is necessary to build on previous efforts and complete them to provide information compliant with Semantic Web principles within agronomic sciences. This adoption would certainly allow the homogenization of multi-scale information, thereby aiding in the discovery of new knowledge. Therefore, we have developed an RDF knowledge-based system, fully compliant with the Semantic Web vision, called Agronomic Linked Data (AgroLD – www.agrold.org) presented hereafter. The aim of our effort is to provide a portal (to discover) and an endpoint (to query) for integrated agronomic information and to aid domain experts in answering relevant biological questions.

The rest of the paper is organized as follows: in the next section, we describe the data sources integrated or used for the integration, the content and architecture of the knowledge-based system. In the following sections, we present the user interface with some examples queries, then we discuss about the contributions and the future directions.

## Materials and Methods

### Information sources

AgroLD was conceived to accommodate molecular and phenotypic information available on various plant species (see Fig 1). The conceptual framework for the knowledge in AgroLD is based on well-established ontologies: GO, SO, PO, Plant Trait Ontology (TO) and Plant Environment Ontology (EO). Among these PO, TO and EO are currently developed by the Planteome project [29] (http://planteome.org). Furthermore, considering the scope of the effort, we decided to build AgroLD in phases. The current phase (phase I) covers information on genes, proteins, ontology associations, homology predictions, metabolic pathways, plant traits, and germplasm, relevant to the selected species. At this stage, we have incorporated the corresponding information from various databases, such as Gramene [30], UniprotKB [31], Gene Ontology Annotation [32], TropGeneDB [33], OryGenesDB [34], Oryza Tag Line [35], GreenPhylDB [36] and SNiPlay [37]. The selection of these data sources was considered based on popularity among domain experts such as GOA, Gramene, and complementary information hosted by the local research community, for instance, Oryza Tag Line and GreenPhylDB. Information on the integrated databases can be found in the documentation page (http://www.agrold.org/documentation.jsp). Table 1 provides a break-down of the data sources and the species covered.

**Table 1.**
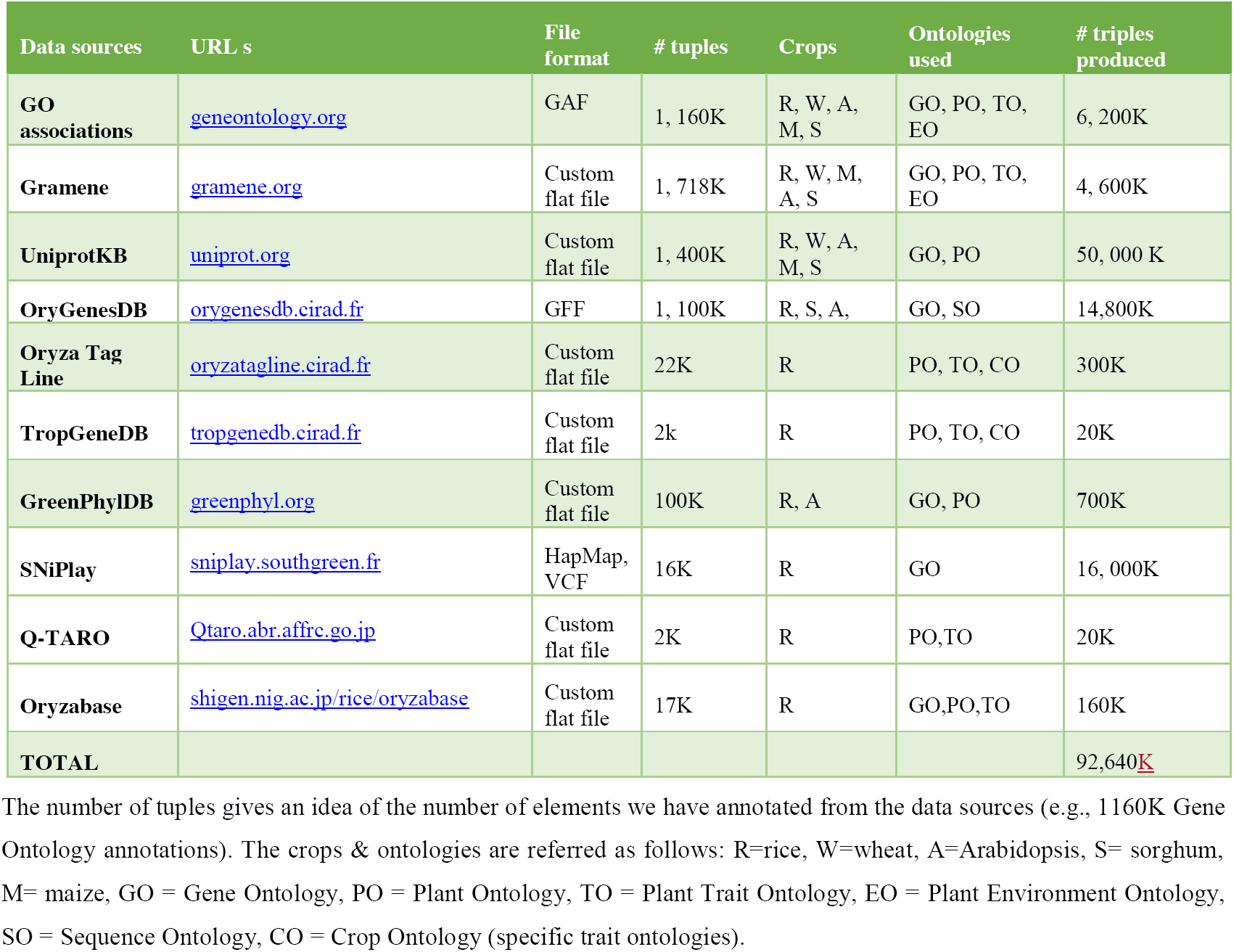
Plant species and data sources in AgroLD.

**Fig 1.**
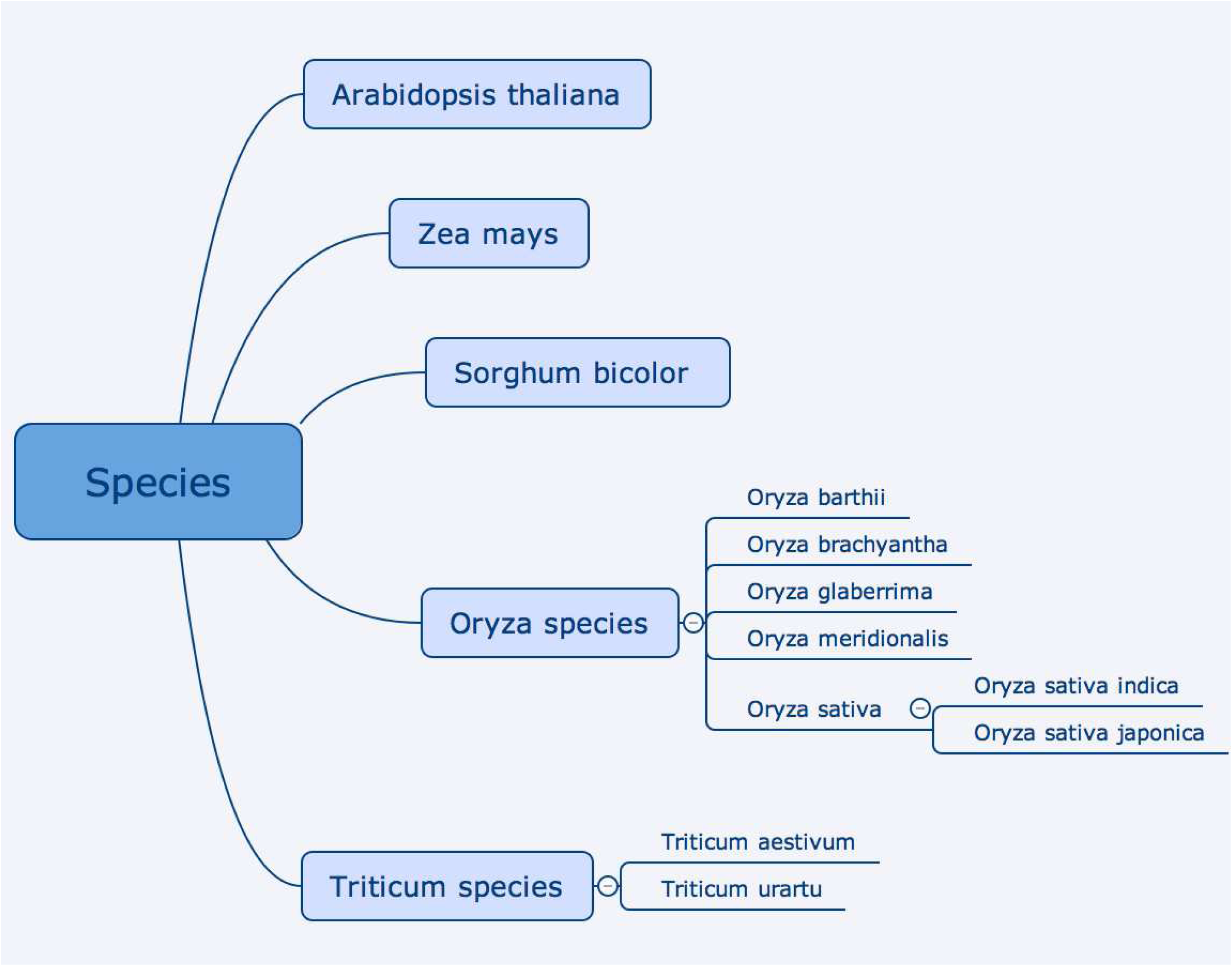
Current plant species included in AgroLD.

### Architecture

AgroLD relies on the RDF and SPARQL technologies for information modelling and retrieval. We use OpenLink Virtuoso (version 7.2) to store and access the RDF graphs. The data from the selected databases were parsed and converted into RDF using a semi-automated pipeline. The pipeline consists of several parsers to handle data in a variety of formats, such as the Gene Ontology Annotation File (GAF) [38], Generic File Format (GFF3) [39], HapMap [40] and Variant Call Format (VCF) [41]. Fig. 2 shows the Extraction-Transform-Load (ETL) processes developed to transform in RDF various source data formats. The source code of the ETL workflow is available on GitHub^12^.

**Fig 2.**
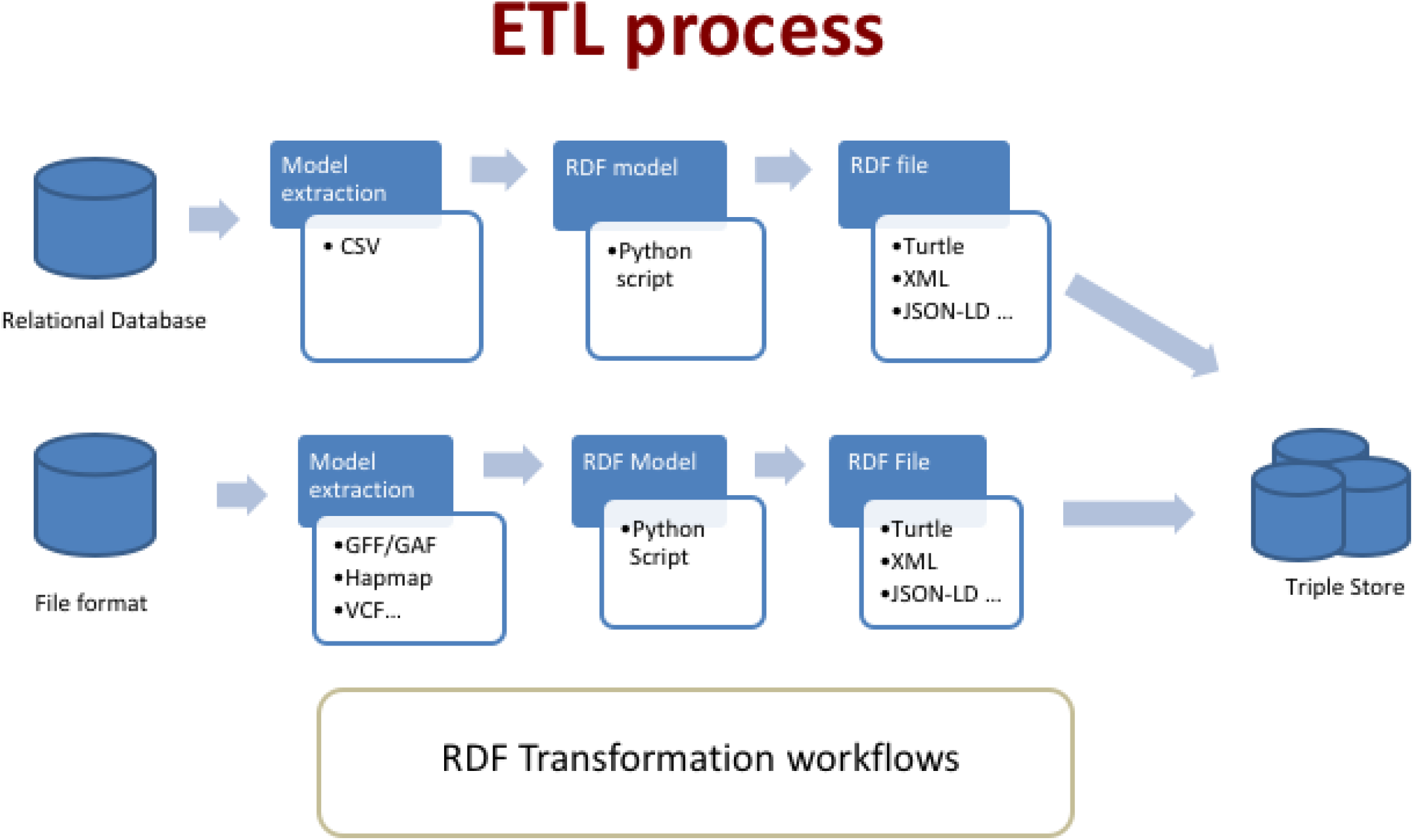
ETL workflow for the various datasets and data formats. The workflow shows two types of process: 1) from relational databases through a CVS file export: in that case, the transformation is tailored for the database model with some Python scripts converters. 2) from standards file formats: in that case, the transformation is generic with some Python packages used as converter tools. The workflow outputs can be produce in various type of RDF format such as turtle, JSON-LD, XML

For this phase, each dataset was downloaded from curated sources and was annotated with ontology terms URIs by reusing the ontology fields when provided by the original source. Additionally, we used the AgroPortal web service API to retrieve the URI corresponding to the taxon available for some data standards such as GFF. At the end of phase 1, early 2018, the AgroLD knowledge base contains around 100 million RDF triples created by converting more than 50 datasets from 10 data sources. Additionally, when available, we used some semantic annotation already present in the datasets such as, for instances, genes or traits annotated respectively with GO or TO identifiers. In that case, we produced additional properties with the corresponding ontologies thus adding 22% additional triples validated manually (see details in Table 1). The OWL versions of the candidate ontologies were directly loaded into the knowledge base but their triples are not counted in the total. We provided in the supplementary file S1 Table, a more comprehensive statistics analysis such as number of triples, classes, entities and properties for each graph stored in the knowledge base.

The RDF graphs are named after the corresponding data sources (protein/qtl ontology annotations being the exception), sharing a common namespace: “http://www.southgreen.fr/agrold/”. The entities in the RDF graphs are linked by shared common URIs. As a design principle, we have used URI schemes made available by the sources (e.g., UniprotKB) or by Identifiers.org registry (http://identifiers.org - [42]). For instances, proteins from UnitProtKB are identified by the base URI: *http://purl.uniprot.org/uniprot/*; genes incorporated from Gramene/Ensembl plants are identified by the base URI: *http://identifiers.org/ensembl.plant/*. New URIs were minted when not provided by the sources or the by Identifiers.org such as TropGene and OryGenesDB; in such cases the URIs take the form ****http://www.southgreen.fr/agrold/*[resource_namespace]/[identifier]***. Furthermore, properties linking the entities took the form: ****http://www.southgreen.fr/agrold/vocabulary/*[property]***. An outline of how the RDF graphs are linked is shown in Fig 3. About entity linking, we used the “key-based approach” which is the most common one. It combines the unique identifier/accession number of the entity shared with the community, with the URI basis pattern of the resource. Moreover, we also respected the “common URI approach” which recommends to use the same URI pattern when the same accession number is used in different datasets. Therefore, defining the same URI for identical entities (represented by identifiers) in different datasets makes it possible to aggregate additional information for this entity. Additionally, we used cross-reference links (represented by identifiers from external datasets) by transforming them into URIs and linked the resource with the predicate “has_dbxref’. This greatly increases the number of outbound links, making AgroLD more integrated with other Linked Open Data. In the future, we will implement a “similarity-based approach” to identify correspondences between entities which have different URIs.

**Fig 3.**
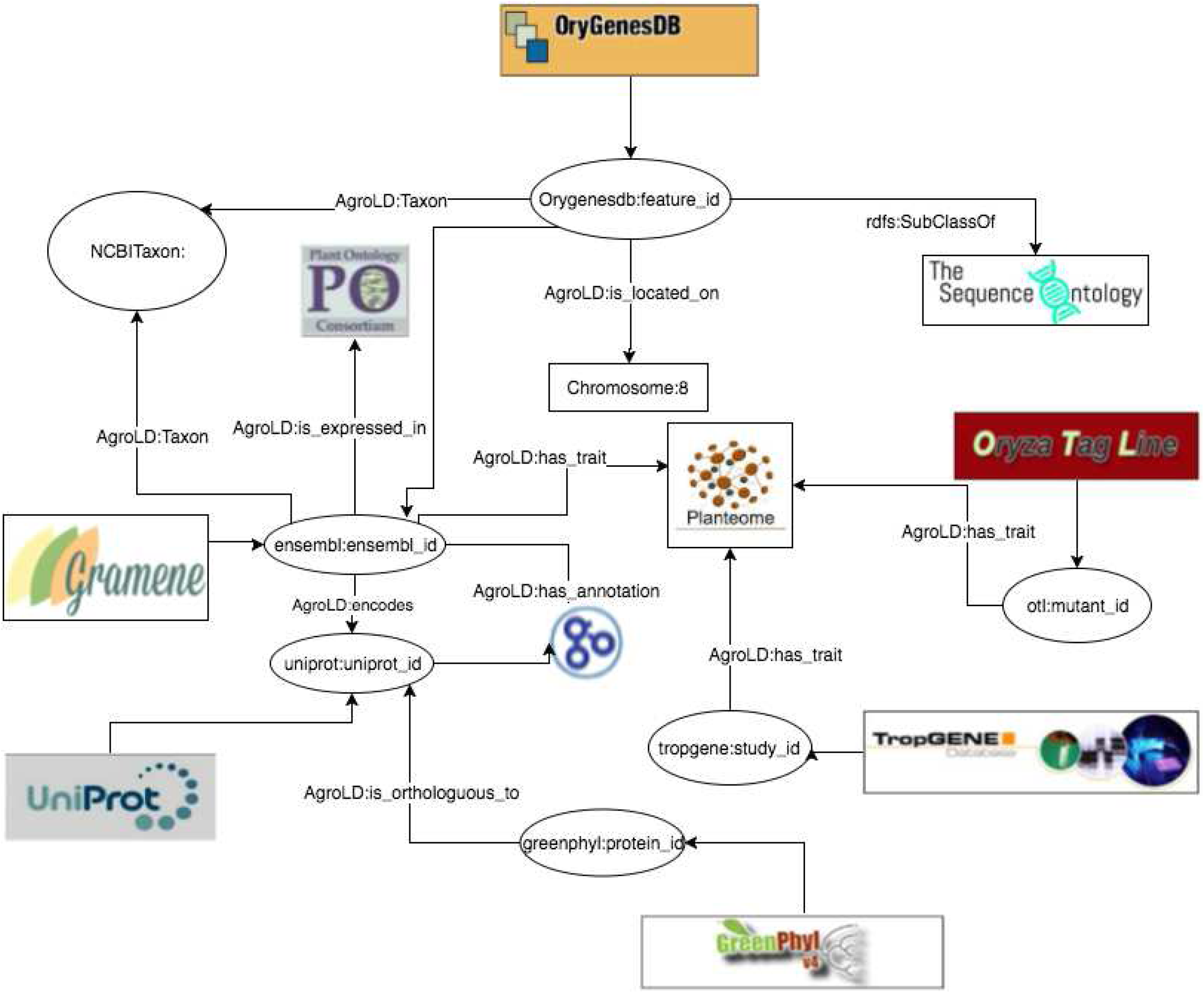
Linking information in AgroLD. The figure illustrates the linking of varies information in AgroLD.

To map the various data types and properties, we developed a lightweight schema (cf. https://github.com/SouthGreenPlatform/AgroLD) that glues classes and properties identified in AgroLD and the corresponding external ontologies. For instance, the class Protein (*http://www.southgreen.fr/agrold/resource/Protein*) is mapped as *owl:equivalentClass* to class polypeptide (*http://purl.obolibrary.org/obo/SO_0000104*) from SO. Similar mappings have been made for properties, e.g., proteins/genes are linked to GO molecular function by the property *http://www.southgreen.fr/agrold/vocabulary/has_function*, which is mapped as *owl:equivalentProperty* to the corresponding Basic Formal Ontology (BFO) term (*http://purl.obolibrary.org/obo/BFO_0000085*). When an equivalent property did not exist, we mapped then to the closest upper level property using *rdfs:subPropertyOf* e.g., the property *has_trait* (*http://www.southgreen.fr/agrold/vocabulary/has_trait*), links proteins to TO terms. It is mapped to a more generic property, *causally related to* in the Relations Ontology [42]. For now, 55 mappings were identified. Furthermore, mappings are both stored side by side with ontologies in AgroPortal, which allows direct links between classes and instances of these classes in AgroLD. For example, the following link will show the external mappings for SO:0000104 (polypeptide) stored in AgroPortal: http://agroportal.lirmm.fr/ontologies/SO/?p=classes&conceptid=http%3A%2F%2Fpurl.obolibrary.org%2Fobo%2FSO_0000104&jump_to_nav=true#mappings. Additionally, classes, properties and resources (e.g., http://www.southgreen.fr/agrold/page/biocyc.pathway/CALVIN-PWY) are dereferenced on a dedicated Pubby server [45]. For details on the graphs, URIs and properties, the reader may refer to AgroLD’s documentation (http://www.agrold.org/documentation.jsp).

### User Interface

The AgroLD platform provides four entry points to access the knowledge base:

- *Quick Search (*http://www.agrold.org/quicksearch.jsp), a faceted search plugin made available by Virtuoso, that allows users to search by keywords and browse the AgroLD’s content;
- *SPARQL Query Editor (*http://www.agrold.org/sparqleditor.jsp), that provides an interactive environment to formulate SPARQL queries;
- *Explore Relationships* visualizer (http://www.agrold.org/relfinder.jsp), which is an implementation of RelFinder [46] that allows users to explore and visualize existing relationships between entities;
- *Advanced Search* (http://www.agrold.org/advancedSearch.jsp), a query form providing entity (e.g., gene) specific information retrieval.

Alternatively, some user management features have been implemented on the platform. Users have the opportunity to save their search and results on a persistent history session attached to their own account. Furthermore, they can manage search history by editing, deleting or re-running previous searches and exporting results according several formats. In the future, we plan to develop some recommendation features and sharing results between users. More detailed descriptions and figures of the different user interfaces will be provided in the following section. Furthermore, other examples are shown in the User Guide available in the supporting information S1 File.

## Results and Discussion

RDF knowledge bases are accessed via SPARQL endpoints and in certain cases equipped with faceted browser interfaces. Using SPARQL endpoints require a minimal knowledge of SPARQL, this may result in the resources not being exploited completely. Alternatively, faceted browser interfaces help the user in getting acquainted with information in the resource (e.g., retrieving a local neighborhood for a particular term), the presence non-textual details (e.g., URIs) in the results could be confusing. To this end, we attempted to lower the usability barrier by providing tools to explore the knowledge base. In this section, we demonstrate the complementary role of the *Advanced Search* and *Explore Relationships* query tools with that of the *SPARQL Query Editor*.

We developed the SPARQL Query Editor based on the YASQE and YASR tools [47] and customized it for our system. The SPARQL language is a powerful tool to mine and extract meaningful information from the knowledge base. In the first example of the supplementary S3 file, we compare two queries to answer the question: “Identify wheat proteins that are involved in root development.”. While the first one (S3_Q1) using a simple search—which is a direct translation of SQL— with the corresponding id (“GO_0048364”, “GO_2000280”) shows 73 entries, the second one (S3_Q2) using a property path query (i.e., query the descending class hierarchy for a given trait ontology term) shows 137 entries, thus more than 80% of additional results. In that case, the use of property path algorithm shows the efficiency in retrieving a comprehensive answer. But the SPARQL language performs also very well with complex queries such as: “Retrieve individuals which have positive SNP variant effect identified for proteins associated with a QTL” available in S3_Q3. This type of query involves several datasets and uses graph traversal property of SPARQL to perform the query.

Because SPARQL is hard to handle for non-technical users, the *SPARQL Query Editor* includes a list of modularized example queries, customizable according to the users’ needs.

For the comparison, we consider a sample question: ‘*Retrieving genes that participate in Calvin cycle*’; (Q6 in the online list of modularized queries). As illustrated in Fig 4, the user can run the query to retrieve the list of genes participating in the given pathway (Fig 4a). Additional information on a gene of interest can be retrieved by clicking on the URI. For example, clicking on AT1GI870 (http://identifiers.org/ensembl.plant/AT1G18270) redirects the users to the gene information provided by Gramene/Ensembl Plants resource (Fig 4b). The query can be saved and the results can be downloaded in a variety of formats such as JSON, TSV, and RDF/XML. Additionally, user defined queries could also be uploaded.

**Fig 4.**
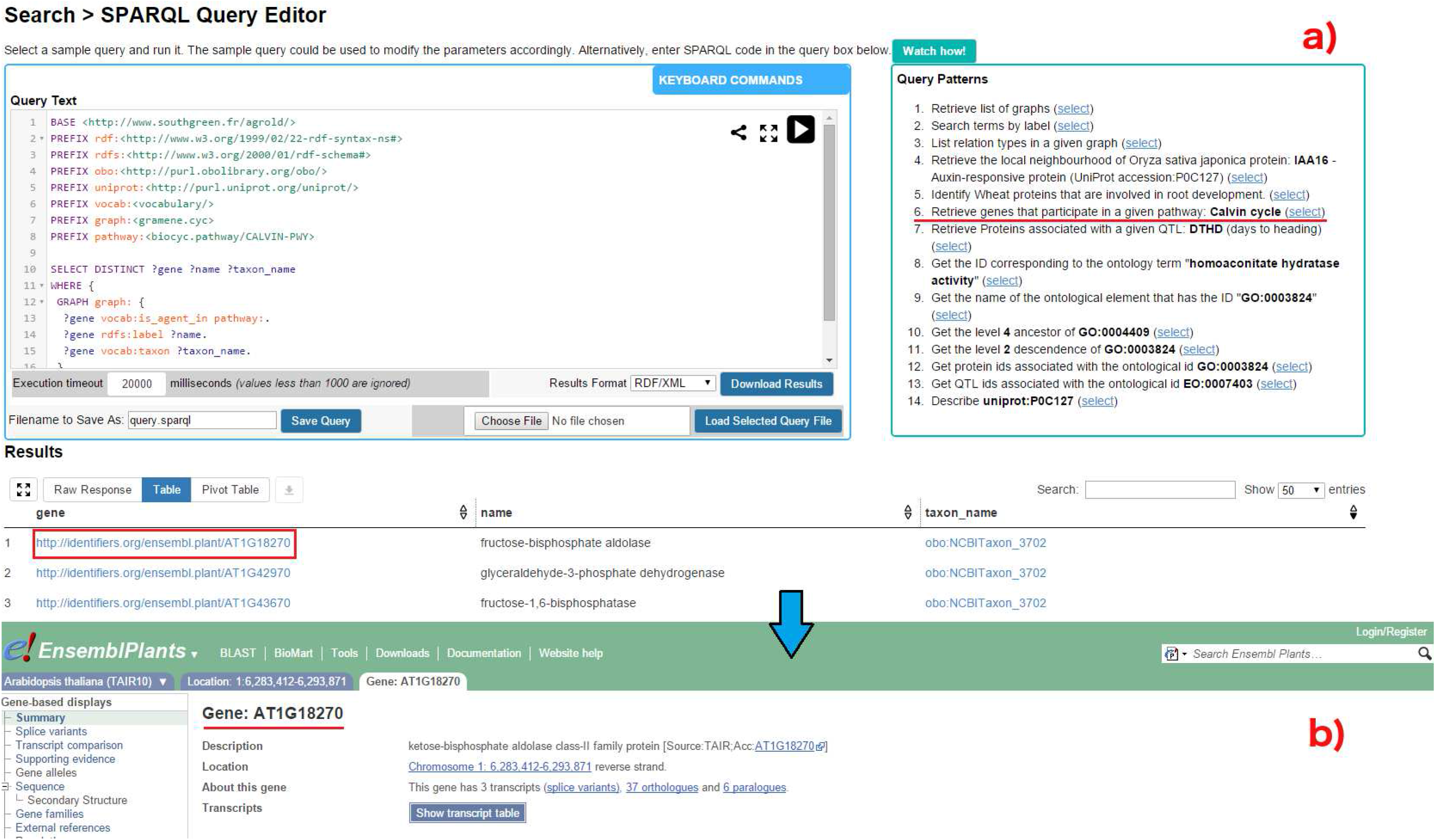
SPARQL Query Editor. Figure illustrates the execution of query Q6: (a) Q6 is one the examples queries on the top-right corner (highlighted in red). On executing the query, the results are rendered below the editor; (b) the user can look up specific genes of interest by clicking on the corresponding URI, which points to the original information source (in this case EsemblPlants).

The *Explore Relationships* tool is based on RelFinder visualization module. This tool aids in visualizing relationships between entities and searching entities by keyword when their URIs are ignored. However, the original version of RelFinder was developed (in ActionScript) and configured for DBpedia. We proposed a configuration and modification of the system suitable for AgroLD. The configuration mainly concerns the SPARQL access point, the properties to be considered for the search of entities and for the description of the resources. Furthermore, we have added some biological examples to guide users. In Fig 5, the tool is used to search for genes involved in Calvin cycle by entering the name of the entities.

**Fig 5.**
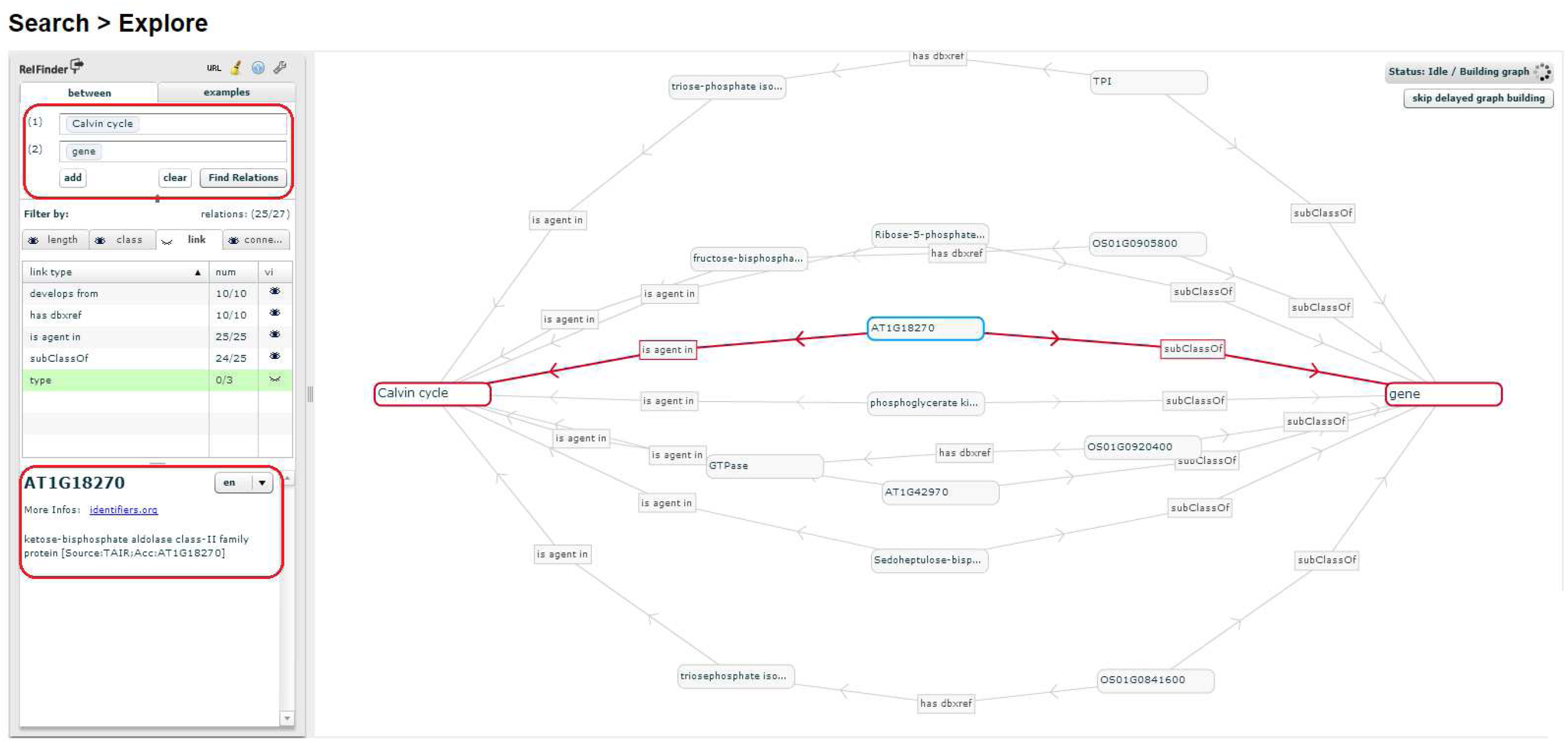
Exploring entity relationships in AgroLD. Figure illustrates differently the results obtained for Q6 using Explore Relationships tool. The results of Q6 can be visualized by entering the concepts (Calvin cycle and gene) in the left panel. On executing the query, all the genes involved in the chosen pathway are revealed. The visualized graph can be altered based on the user interest. Additionally, a gene could be selected (circled on the left) and further explored by clicking on the *More Info* link which directs the user to the information source

The *Advanced Search* query form is based on the REST API suite (http://www.agrold.org/api-doc.jsp), developed completely within the AgroLD project. The aim of this feature is to provide non-technical users with a tool to query the knowledge base while hiding the technical aspects of SPARQL query formulation. Fig 6 illustrates steps involved in retrieving information for Q6, using the query form:

**Fig 6.**
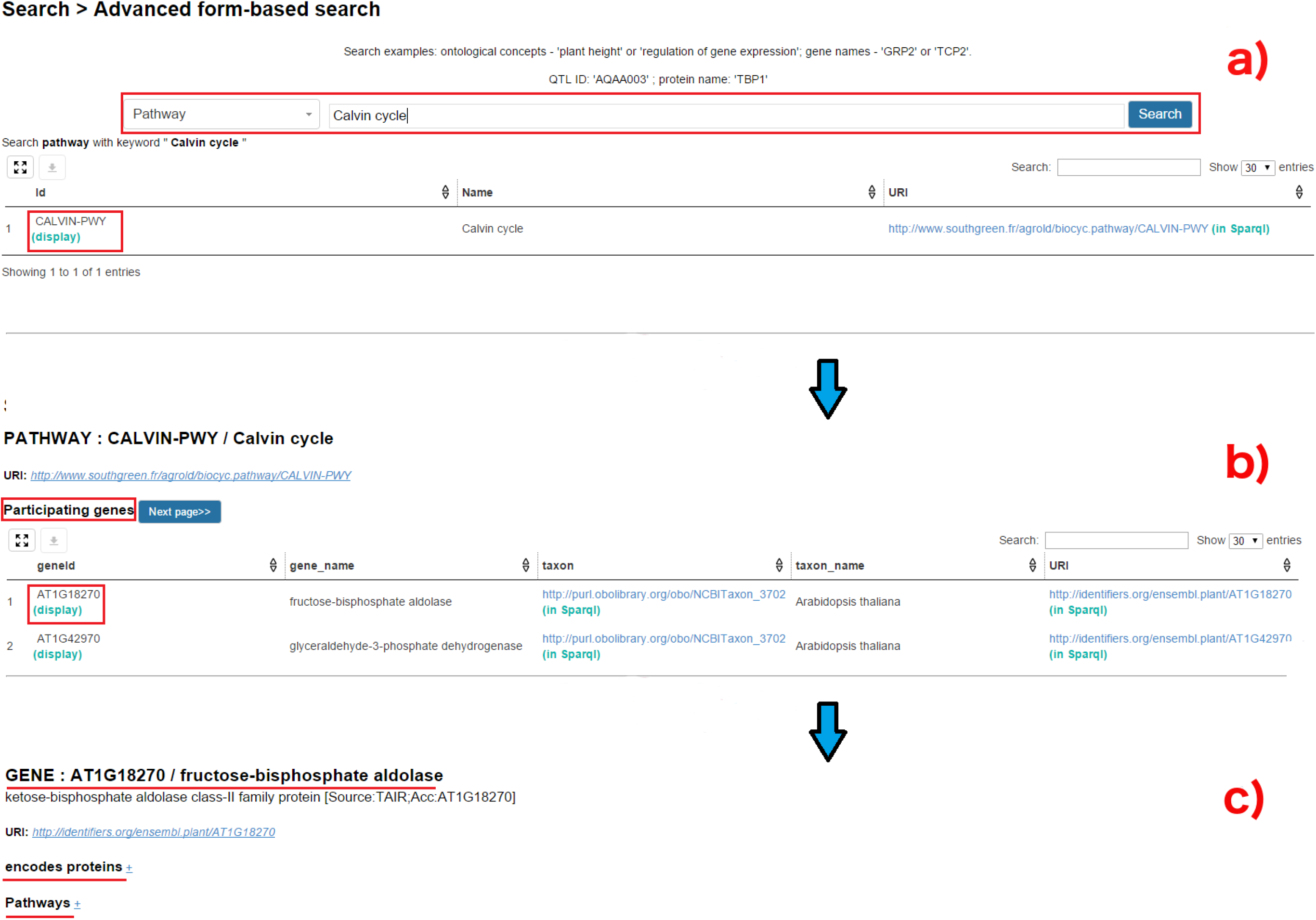
Advanced Search query form: Figure demonstrates the steps involved in retrieving the results for Q6 using the Advanced Search query form: (a) query Q6 can be executed by selecting the type of entity (Pathways – highlighted in red) to search and entering the name of the entity (Calvin cycle). The API then displays the matched results; (b) Clicking on the result displays the genes participating in Calvin cycle; (c) selecting a gene of interest displays more information pertaining to that gene, for instance, encoding proteins and pathways this selected gene participates in.

a. The user selects *Pathways* from the list of entities and enters the pathway of interest, in this case, Calvin cycle (Fig 6a);
b. The list of genes involved in the pathway can be retrieved by selecting the pathway.

Furthermore, information on a gene of interest can be retrieved by selecting the specific gene (Fig 6b). For instance, clicking on AT1GI870 (Fig 6c) displays all the proteins the gene encodes and the pathways the gene participates in (apart from Calvin cycle). The RESTful API supports the query form and was developed for programmatic retrieval of entity specific knowledge represented in AgroLD. The current version of the API suite (ver. 1) can be used to retrieve gene and protein information, metabolic pathways, and proteins associated with ontological terms. This is achieved by querying entity by name or identifier.

### User Evaluation

AgroLD is being actively developed based on usability testing sessions conducted with domain experts, including doctoral students in biology, curators and senior researchers. Test sessions were designed to measure if:

- Resources integrated in AgroLD are useful;
- AgroLD is easy to use.

For the evaluation of semantic search systems, Elbedweihy et al. [48] recommend a survey of users based on their experience with a few queries submitted to the system. We have used this approach to collect user opinions, comments and suggestions via a feedback form directly within the AgroLD web application. The form includes some questions from the “System Usability Scale” questionnaires [49] and other questions that we considered important. The three main criteria evaluated are:

1. Usability –ease to submit a query (number of attempts, time required) and presentation of the results;
2. Expressiveness – type of queries a user is able to formulate (e.g., keywords or more complex expressions);
3. Performance –speed, correctness and completeness of the results.

Recently, 20 participants were invited during 3 testing sessions, to search for concepts, genes, or pathways of their interests; and the online form was active (http://agrold.org/survey.jsp) to allow new feedbacks during the exploitation phase. Each question had 5 possible answers ranked from the highest to the lowest note (5 to 1). We reported the results of these sessions in S2 File as a supplementary document.

Globally, participants found the platform useful and easy to use. Overall, the idea of data navigation and traversal through knowledge graphs was well received. However, many of them needed help with some features. The general observation is that testing users ranked *Advanced Search* first then *Quick Search* after. We explain this by the display output that looks friendlier for Advanced Search. *Quick Search* won votes for usability and performance despite several comments to improve the ranking and presentation of results (4 user’s comments). *Advanced and Explore search* got average scores but good comments on the capability of discovering unexpected results (e.g., nearest neighbour entities in the graph for the Explore Search and additional results from external Web services for Advanced Search). With no surprise, evaluation results show the *SPARQL Query Editor* is the most difficult to handle. We mitigate this by offering examples of query pattern to help users handle query formulation. In the future, we will improve the examples by offering a large spectrum of search type which will follow the new phase of data integration. Furthermore, we will provide links to some SPARQL tutorials in the documentation. These user feedbacks reinforced the need for knowledge bases such as AgroLD, wherein users could retrieve information across various data types and sources. This knowledge discovery is supported by the use of shared URI schemes and domain ontologies. The testing sessions also helped us to identify areas for further improvement. Plus, we received suggestions on improving the AgroLD’s coverage with more data types such as gene expression data, and protein-protein interactions. Considering, linked data and Semantic Web are still not widely adopted in agronomy, increasing AgroLD’s coverage will be an incremental process engaging our user community. This situation is expected to improve with new community efforts such as the Agrisemantics RDA Working Group (https://rd-alliance.org/groups/agrisemantics-wg.html), which role is to reinforce the adoption of semantic technologies in the agri-food domain. We may also mention the AgBioData consortium (https://www.agbiodata.org, [2]) which promotes the FAIR (Findable, Accessible, Interoperable and Reusable) data principles [50] within agricultural research.

Furthermore, we observed that although the information integrated in AgroLD came from curated sources, scientists often prefer to validate these knowledge statements against assertions made in scientific articles. Currently, we have implemented an external Web Services as part of the *Advanced Search Form* to automatically search for publications related to a protein or gene of interest in PubMed Central and aggregates them within the result of the AgroLD query. However, this feature does not provide detailed (sentence level) assertions described in those publications. This is an area that requires further work. With the recent developments towards making text mined (sentence level) annotations available as RDF [51], query federation can be explored to retrieve entity specific assertions. This would serve as an additional provenance layer.

### Limits and Perspectives

With the achievement of the first phase of AgroLD, many plant scientists can benefit from the interoperability of the data, but user feedback reveals some limitations and challenges on the current version of AgroLD. In order to achieve the expectations of the scientists for the use of Semantic Web technologies in agronomy, a number of issues need to be addressed:

- The coverage content has to be extended to a larger number of biological entities (e.g., miRNA, mRNA) or interaction between them (e.g., co-expression, regulation and interaction networks) in order to capture a broad view of the molecular interactions.
- We have observed many information remains hidden in RDF literal contents such as biological entities or relationship between them. This information is poorly annotated (i.e., plain text not formally expressed) and new research methods to identify biological entities and reconstruct their relations further allowing the discovery of relevant links between related resources are required.
- The explosion of data in agronomy forces database providers to augment the frequency of their releases. The survey shows a growing interest of using up to date information from the original sources. This have to be taken into account for the updating process in AgroLD.
- The user interfaces show some limitations to manage responses with large number of results, e.g., to filter and rank them with precision score.

These limitations identified in the current version of AgroLD will be improved in the following versions. We will focus on the following areas:

- User Interface: we plan to explore features offered by Elastic search tool (https://www.elastic.co), to enabling *Quick Search* retrieving more textual information and hiding the technical details. Further, we will improve the performance and expand the API suite to cover other entities represented in AgroLD (e.g., genomic annotation and homology information).
- Content: integrate information on gene expression such as IC4R [52], Gene Expression Atlas [53], on gene regulatory networks such as RiceNetDB [54] and explore linking text-mined annotations from publications. Support molecular interaction networks per species and also allow knowledge transfer between species.
- Knowledge discovery: explore methods to aid generating hypotheses by retrieving implicit knowledge, e.g., inference rules, automatic data linking, entity recognition, text mining, automatic semantic annotations.
- Data provenance: develop a provenance and annotation model. Set up a validation process to allow users validating computed facts such as semantic annotations automatically produced and attached to a biological entity.
- Updates: To keep AgroLD updated with the latest available data, by processing regular data updates and potentially re-building the entire repository from scratch every 12 months.^3^ Additionally, we plan to fully automate the current ETL workflow.

## Conclusion

Data in the agronomic domain are highly heterogeneous and dispersed. For agronomic researchers to make informed decisions in their daily work it is critical to integrate information at different scales. Current traditional information systems are not able to exploit such data (i.e., genes, proteins, metabolic pathways, plant traits, and phenotypes), in efficient way. To this end, the application of Semantic Web, initiated in the biomedical domain, provides a good example to follow by capitalizing on previous experiences and addressing weaknesses.

To further build on this line of research in agronomy, we have developed AgroLD. We have demonstrated the advantages of AgroLD in data integration over multiple data sources using plant domain ontologies and Semantic Web technologies. To date, AgroLD contains 100M of triples created by transforming more than 50 datasets coming from 10 data and annotating with 10 ontologies. The impact of AgroLD is expected to grow with an increase in coverage (with respect to the species and the data sources) and user inputs. For instance, when user feedback and implementation of inference rules are put within a context that supports searching and recommendations, then we have the beginnings of a platform that can support automated hypotheses generation.

AgroLD is one of the first RDF linked open data knowledge-based system in the agronomic domain. It demonstrates a first step toward adopting the Semantic Web technologies to facilitate research by integrating numerous heterogeneous data and transforming them into explicitly knowledge thanks to ontologies. We expect AgroLD will facilitate the formulation of new scientific hypotheses to be validated with its knowledge-oriented approach.

## Funding

This research was supported by the Computational Biology Institute of Montpellier (ANR-11-BINF-0002), the Institut Francais de Bioinformatique (ANR-11-INBS-0013), the Labex Agro (ANR-10-LABX-001-01) all bypass of the French ANR *Investissements d’Avenir* program.

## Authors’ contributions

AV designed and implemented the AgroLD project and wrote the manuscript. GT designed and implemented the API and the website. NEH contributed to the integration of data and set up of the RDF store. IC tested and formulated biological queries. VG contributed to the integration of data. CJ reviewed the manuscript. MR helped conceive the AgroLD project and reviewed the manuscript. PL conceived, designed, implemented the AgroLD project and wrote the manuscript. All the authors approved the final manuscript.

## Acknowledgments

Authors thank the technical staffs of the South Green Bioinformatics platform for their support. Authors thank the providers of databases listed in Fig 1, who kindly gave access to their publicly datasets. Authors thank the expert biologists and bioinformaticians who contributed to the testing sessions and helped us to improve the content of the system and the user interface. Authors specially thank Dr. Patrick Valduriez and Dr. Eric Rivals for their supports and advises in this project.

## Supporting information

**S1 File. AgroLD User Guide.** This document shows how to use the various features of the platform.

**S1 Table. AgroLD graph statistics.**

**S2 File. Report of the online survey.** Report of 3 sessions evaluating the AgroLD user interfaces.

**S3 File. Examples of SPARQL queries.** Example of SPARQL queries showing the benefits of property path algorithm, and complex queries.

## Footnotes

^1^https://doi.org/10.5281/zenodo.1294660

^2^https://github.com/SouthGreenPlatform/AgroLD

^3^ Processing regular data update is a hard issue has the original databases do not always provide an automatic way to obtain the differential data between releases. From experience, we know that regularly rebuilding the entire knowledge base is for us a good alternative to avoid dealing with data diffs.

